# Tourmaline: a containerized workflow for rapid and iterable amplicon sequence analysis using QIIME 2 and Snakemake

**DOI:** 10.1101/2021.09.15.460495

**Authors:** Luke R. Thompson, Sean R. Anderson, Paul A. Den Uyl, Nastassia V. Patin, Shen Jean Lim, Grant Sanderson, Kelly D. Goodwin

**Affiliations:** Northern Gulf Institute, Mississippi State University, Mississippi State, MS, USA; Ocean Chemistry and Ecosystems Division, Atlantic Oceanographic and Meteorological Laboratory, National Oceanic and Atmospheric Administration, Miami, Florida, USA; Cooperative Institute for Great Lakes Research, University of Michigan, Ann Arbor, MI, USA; Cooperative Institute for Marine and Atmospheric Studies, Rosenstiel School of Marine and Atmospheric Science, University of Miami, Miami, FL, USA; Marine Science Department, University of Hawaii, Hilo, HI, USA

**Keywords:** amplicon sequencing, metabarcoding, environmental DNA, eDNA, microbiome, meta-analysis

## Abstract

**Background:** Amplicon sequencing (metabarcoding) is a common method to survey diversity of environmental communities whereby a single genetic locus is amplified and sequenced from the DNA of whole or partial organisms, organismal traces (e.g., skin, mucus, feces), or microbes in an environmental sample. Several software packages exist for analyzing amplicon data, among which QIIME 2 has emerged as a popular option because of its broad functionality, plugin architecture, provenance tracking, and interactive visualizations. However, each new analysis requires the user to keep track of input and output file names, parameters, and commands; this lack of automation and standardization is inefficient and creates barriers to meta-analysis and sharing of results.

**Findings:** We developed Tourmaline, a Python-based workflow that implements QIIME 2 and is built using the Snakemake workflow management system. Starting from a configuration file that defines parameters and input files—a reference database, a sample metadata file, and a manifest or archive of FASTQ sequences—it uses QIIME 2 to run either the DADA2 or Deblur denoising algorithm, assigns taxonomy to the resulting representative sequences, performs analyses of taxonomic, alpha, and beta diversity, and generates an HTML report summarizing and linking to the output files. Features include support for multiple cores, automatic determination of trimming parameters using quality scores, representative sequence filtering (taxonomy, length, abundance, prevalence, or ID), support for multiple taxonomic classification and sequence alignment methods, outlier detection, and automated initialization of a new analysis using previous settings. The workflow runs natively on Linux and macOS or via a Docker container. We ran Tourmaline on a 16S rRNA amplicon dataset from Lake Erie surface water, showing its utility for parameter optimization and the ability to easily view interactive visualizations through the HTML report, QIIME 2 viewer, and R- and Python-based Jupyter notebooks.

**Conclusions:** Automated workflows like Tourmaline enable rapid analysis of environmental and biomedical amplicon data, decreasing the time from data generation to actionable results. Tourmaline is available for download at github.com/aomlomics/tourmaline.

## Background

Earth’s environments are teeming with environmental DNA (eDNA): free and cellular genetic material from whole microorganisms (Consortium 2012; Thompson *et al*. 2017) or remnants of larger macroorganisms (Deiner *et al*. 2017; Compson *et al*. 2020). This eDNA can be collected, extracted, and sequenced to reveal the identities and functions of the organisms that produced it. Amplicon sequencing (metabarcoding), whereby a short genomic region is amplified and sequenced using polymerase chain reaction (PCR) from an environmental or experimental community’s eDNA, is a popular method for measuring taxonomic diversity of microbiomes and environmental samples (Zaiko *et al*. 2015; Deiner *et al*. 2017; Ruppert, Kline and Rahman 2019). PCR primers have been used to generate amplicons of the bacterial 16S rRNA gene in studies of human and animal-associated microbiota (Turnbaugh *et al*. 2006; Ahn *et al*.2013; Kartzinel *et al*. 2019), as well as environmental microbiota (Sunagawa *et al*. 2015; Thompson *et al*. 2017). Other regions that are commonly targeted include the fungal internal transcribed spacer (ITS) regions between rRNA genes (Abarenkov *et al*. 2010), the 18S rRNA gene of eukaryotes (Vargas *et al*. 2015), the mitochondrial cytochrome oxidase I (COI) gene of invertebrate and vertebrate eDNA (Leray *et al*. 2013), and the mitochondrial 12S rRNA gene of fish (Miya *et al*. 2015). Information gained from amplicon metabarcoding has far reaching implications for human health (e.g., microbiome research), ecosystem function and conservation, and resource management (Thomsen and Willerslev 2015; Halfvarson *et al*. 2017a).

Combining datasets from amplicon surveys and performing meta-analysis can reveal patterns impossible to observe from individual studies and also provide more power in statistical analyses. One example of the power of meta-analysis in amplicon surveys is the Earth Microbiome Project (Thompson *et al*. 2017), which used a single 16S rRNA amplicon method, metadata standard, and denoising algorithm to sequence and analyze over 25,000 microbial communities from around Earth. Standardization of methods is critical for comparability across studies. A lack of standardized methods is one of the main factors limiting cross-comparison of microbiome or eDNA datasets in meta-analyses (Dickie *et al*. 2018; Harper *et al*. 2019). While standardizing detailed laboratory methods across many labs in multiple countries is a major challenge, standardizing analysis methods and metadata formats is much more feasible.

A popular approach for standardizing amplicon data analysis is to develop analysis pipelines or workflows (Reiter *et al*. 2021) to run on local or networked computing resources or in the cloud. Amplicon workflows have become increasingly popular and include pipelines like [Anacapa](https://github.com/limey-bean/Anacapa) (Curd *et al*. 2019), Banzai (https://github.com/jimmyodonnell/banzai), [PEMA](https://github.com/hariszaf/pema) (Zafeiropoulos *et al*. 2020), [Ampliseq](https://github.com/nf-core/ampliseq) (Straub *et al*. 2020), [Cascabel](https://github.com/AlejandroAb/CASCABEL) (Asbun *et al*. 2019), [dadasnake](https://github.com/a-h-b/dadasnake) (Weißbecker, Schnabel and Heintz-Buschart 2020), CoMA (Hupfauf *et al*. 2020), [ASAP 2](https://hts.iit.edu/asap2) (Tian and Imanian 2022), and tagseq (https://github.com/shu251/tagseq-qiime2-snakemake). However, stand-alone pipelines typically do not have access to a wide range of datasets, and many pipelines are bespoke workflows with a jumble of custom scripts for the user to navigate. A variation on this model would be a stand-alone workflow that runs on top of a widely used amplicon software tool, such that it could take advantage of the functionality of the underlying tool and evolve with it, while remaining interoperable with any other analyses using this tool. If deployed on the cloud or in a container, it would also be portable enough to run on larger datasets.

QIIME 2 (Bolyen *et al*. 2019) is a popular software package that provides command-line, Python, and graphical user interfaces for amplicon sequence analysis from raw FASTQ sequences to observation tables, statistical analyses, and interactive visualizations. QIIME 2 supports DADA2 (Callahan *et al*. 2016) and Deblur (Amir *et al*. 2017) plugins for denoising amplicon sequence data. Snakemake (Köster and Rahmann 2012) is a workflow management system that is popular in the bioinformatics community. Snakemake manages input and output files in a defined directory structure, with commands defined in a *Snakefile* as ‘rules,’ and parameters and initial input files set by the user in a configuration file. Snakemake ensures that only the commands required for requested output files not yet generated are run, saving time and computation when re-running part of a workflow.

Here, we present Tourmaline (github.com/aomlomics/tourmaline), an amplicon analysis pipeline that uses Snakemake to run QIIME 2 commands for core analysis and interactive visualization—plus workflow-specific commands that generate an HTML report of output and summary tables and figures of data and metadata—with rapid analysis aided by workflow iterability and scalability, support for multiple cores, a Docker container, and detailed documentation. After cloning the initial Tourmaline directory from GitHub and setting up the input files and parameters, only a few simple shell commands are required to execute the Tourmaline workflow. Outputs are stored in a standard directory structure that is the same for every Tourmaline run, facilitating data exploration and sharing, parameter optimization, and downstream analysis. Because of this standard directory structure, different runs that utilize different parameters (e.g., DADA2 truncation lengths) can be easily compared, facilitated by a helper script that makes a new copy of the Tourmine directory from an existing one. Every Tourmaline run produces an HTML report containing a summary of metadata and outputs, with links to web-viewable QIIME 2 visualization files. QIIME 2 artifact files can be fed directly into Python- and R-based analysis packages. In addition to running natively on Mac and Linux platforms, Tourmaline can be run in any computing environment using Docker containers. In this paper, we describe the Tourmaline workflow, and apply it to a downsampled 16S rRNA gene dataset from surface waters of Lake Erie. The tutorial includes guidance on evaluating output to refine parameters for the workflow and showcases the HTML report, interactive visualizations, and R- and Python-based analysis notebooks for biological insight into amplicon datasets.

## Findings

### Workflow

#### Overview

Tourmaline is a Snakemake-based bioinformatics workflow that operates in a defined directory structure (Fig. 1). Installation involves installing QIIME 2 and other dependencies or installing the Docker container. The starting directory structure is then cloned directly from GitHub and is built out through Snakemake commands, defined as ‘rules’ in *Snakefile*. Tourmaline provides seven high-level ‘pseudo-rules’ for each of DADA2 paired-end, DADA2 single-end, and Deblur (singleend), running denoising and taxonomic and diversity analyses via QIIME 2 and other programs, encompassing commonly used analyses in microbiome and eDNA research. For each type of processing, there are four steps: (1) the *denoise* rule imports FASTQ data and runs denoising, generating a feature table and representative sequences; (2) the *taxonomy* rule assigns taxonomy to representative sequences; (3) the *diversity* rule does representative sequence curation, core diversity analyses, and alpha and beta group significance; and (4) the *report* rule generates an HTML report of the metadata, inputs, outputs, and parameters. Steps 2–4 have two modes each, *unfiltered* and *filtered*, thus making seven pseudo-rules total. The difference between the *unfiltered* and *filtered* commands is that in the *taxonomy_filtered* command, undesired taxonomic groups or individual sequences from the representative sequences and feature table are filtered (removed). The *diversity* and *report* rules are identical for *unfiltered* and *filtered* commands, except the outputs go into separate subdirectories. In addition to the 21 pseudo-rules (3 denoising methods with 7 pseudo-rules each), there are 47 regular rules defined in *Snakefile* that perform the actual QIIME 2, Python, and shell commands of the workflow (Fig. S1).

**Figure 1.**
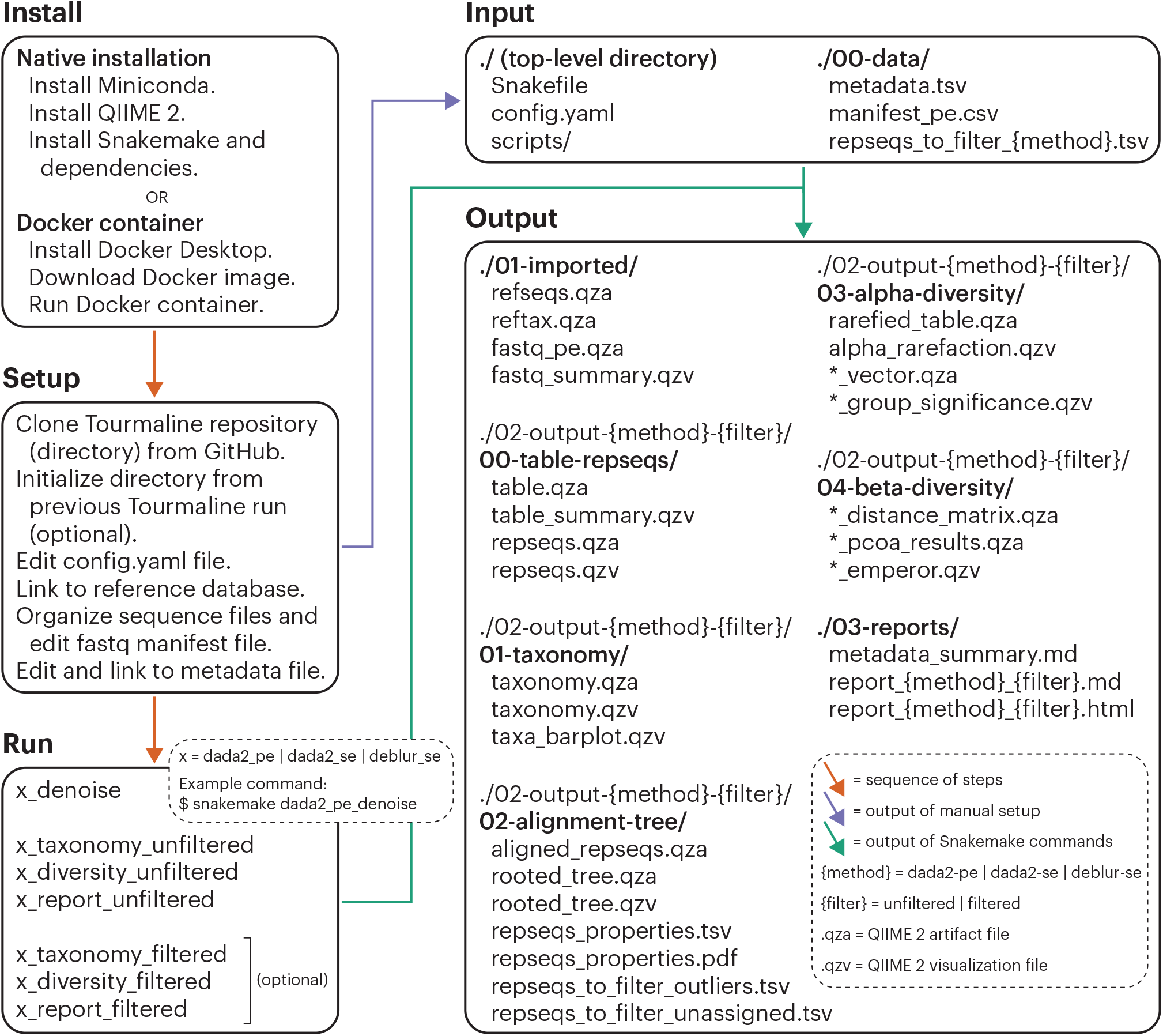
The Tourmaline workflow. Install natively (macOS, Linux) or using a Docker container. Setup by cloning the Tourmaline repository (directory) from GitHub, initializing the directory from a previous run (optional), editing the configuration file (*config.yaml*, Table S1), creating symbolic links to the reference database files, organizing the sequence files and/or editing the FASTQ manifest file, and editing and creating a symbolic link to the metadata file. Run by calling the Snakemake commands for *denoise, taxonomy, diversity*, and *report*—or running just the *report* command to generate all output if the parameters do not need to be changed between individual commands. It is recommended but not required to run the *unfiltered* commands before the *filtered* commands. The primary input and output files are listed. Detailed instructions for each step are provided in the Tourmaline Wiki (github.com/aomlomics/tourmaline/wiki).

#### Test dataset

Tourmaline comes with a test dataset of 16S rRNA gene (bacteria/archaea) amplicon data from surface waters of Western Lake Erie in summer 2018 (see Methods). The sequence data were subsampled to 1000 sequences per sample to allow the entire workflow to run in ~10 minutes. This test dataset is used throughout the paper to demonstrate the capabilities of Tourmaline.

#### Documentation

Full instructions for using the Tourmaline workflow, including installation, cloning, and editing the config file, are described in the Tourmaline Wiki at github.com/aomlomics/tourmaline/wiki. Some experience with the command line, QIIME 2, and Snakemake is helpful to use Tourmaline; basic tutorials for each of these are provided at github.com/aomlomics/tutorials.

#### Installation

The workflow requires QIIME 2 (version 2021.2) plus several dependencies, which can be installed natively in a Conda environment (instructions on GitHub) or via a Docker container using the Docker image from [DockerHub](https://hub.docker.com/repository/docker/aomlomics/tourmaline). Tourmaline is installed by cloning the GitHub repository to the current directory with *git clone* https://github.com/aomlomics/tourmaline. This step is repeated any time a new iteration of Tourmaline is needed, and new copies can be initialized using a helper script (described below).

#### Snakefile

As a Snakemake workflow, Tourmaline has as its core files (1) a *Snakefile* that provides all the commands (rules) that comprise the workflow and (2) a *config.yaml* file that provides the input files and parameters for the workflow. *Snakefile* contains all of the commands used by Tourmaline, which invoke QIIME 2 commands, helper scripts (see below), or generate output directly. The main analysis features and options supported by Tourmaline, as specified in *Snakefile*, are as follows:

- FASTQ sequence import using a manifest file, or use a pre-imported FASTQ .qza file.
- Denoising with DADA2 (Callahan *et al*. 2016) (paired-end and single-end) and Deblur (Amir *et al*. 2017) (single-end).
- Feature classification (taxonomic assignment) with options of naive Bayes (Bokulich *et al*. 2018), consensus BLAST (Camacho *et al*. 2008), and consensus VSEARCH (Rognes *et al*.2016).
- Feature filtering by taxonomy, sequence length, feature ID, and abundance/prevalence.
- De novo multiple sequence alignment with MUSCLE (Edgar 2004), Clustal Omega (Sievers and Higgins 2014), or MAFFT (Katoh and Standley 2013) (with masking) and tree building with FastTree (Price, Dehal and Arkin 2009).
- Outlier detection with odseq (Jehl, Sievers and Higgins 2015).
- Interactive taxonomy barplot.
- Tree visualization using Empress (Cantrell *et al*. 2021).
- Alpha diversity, alpha rarefaction, and alpha group significance with four metrics: number of observed features, Faith’s phylogenetic diversity, Shannon diversity, and Pielou’s evenness.
- Beta diversity distances, principal coordinates, Emperor (Vázquez-Baeza *et al*. 2013) plots, and beta group significance (one metadata column) with four metrics: unweighted and weighted UniFrac (Lozupone *et al*. 2010), Jaccard distance, and Bray–Curtis distance.
- Robust Aitchison PCA and biplot ordination using DEICODE (Martino *et al*. 2019).

#### Config file

The configuration file *config.yaml* includes paths to input files and parameters for QIIME 2 commands and other steps. Default settings have been chosen to balance run performance and accuracy and to work with the test data. For user data, all parameters should be checked and possibly adjusted for appropriateness with the dataset; see Table S1, Fig. 1, and the Wiki section *Setup* for guidance.

#### Input files

Tourmaline requires three categories of input files: (1) Reference database: a FASTA file of reference sequences (*refseqs.fna*) and a tab-delimited file of taxonomy (*reftax.tsv*) for those sequences, or their imported QIIME 2 artifact equivalents (*refseqs.qza*, *reftax.qza*); (2) Amplicon data: demultiplexed FASTQ sequence files and FASTQ manifest file(s) (*manifest_pe.csv*, *manifest_se.csv*) mapping sample names to the location of the sequence files, or their imported QIIME 2 equivalents (*fastq_pe.qza*, *fastq_se.qza*); and (3) Metadata: a tab-delimited sample metadata file (*metadata.tsv*) with sample names in the first column matching those in the FASTQ manifest file. See the Wiki section *Setup* for guidance on input file paths and use of symbolic links to avoid storing multiple copies of large input files.

#### Run the workflow

The workflow is run using Snakemake commands. For example, if using DADA2 paired-end method without any filtering (see below), the commands would be (1) *snakemake dada2_pe_denoise*, (2) *snakemake dada2_pe_taxonomy_unfiltered*, (3) *snakemake dada2_pe_diversity_unfiltered*, and (4) *snakemake dada2_pe_report_unfiltered*. Alternatively, the entire workflow can be run at once with the last command, *snakemake dada2_pe_report_unfiltered*.

### Outputs

The outputs of each step of Tourmaline are described following a test run with the Lake Erie test data that comes with the GitHub repository. For each command, the main parameters used and list of output files generated in those commands are provided (Fig. 2). Accompanying the list of output files is guidance for evaluating them to choose parameters for subsequent steps (Fig. 2), with screenshots of the Tourmaline-specific output files (Fig. 3) and both QIIME 2 and Tourmaline-specific output files (Fig. S3). A video version of the tutorial is also available on YouTube (https://youtu.be/1lFyeswXSX8).

**Figure 2.**
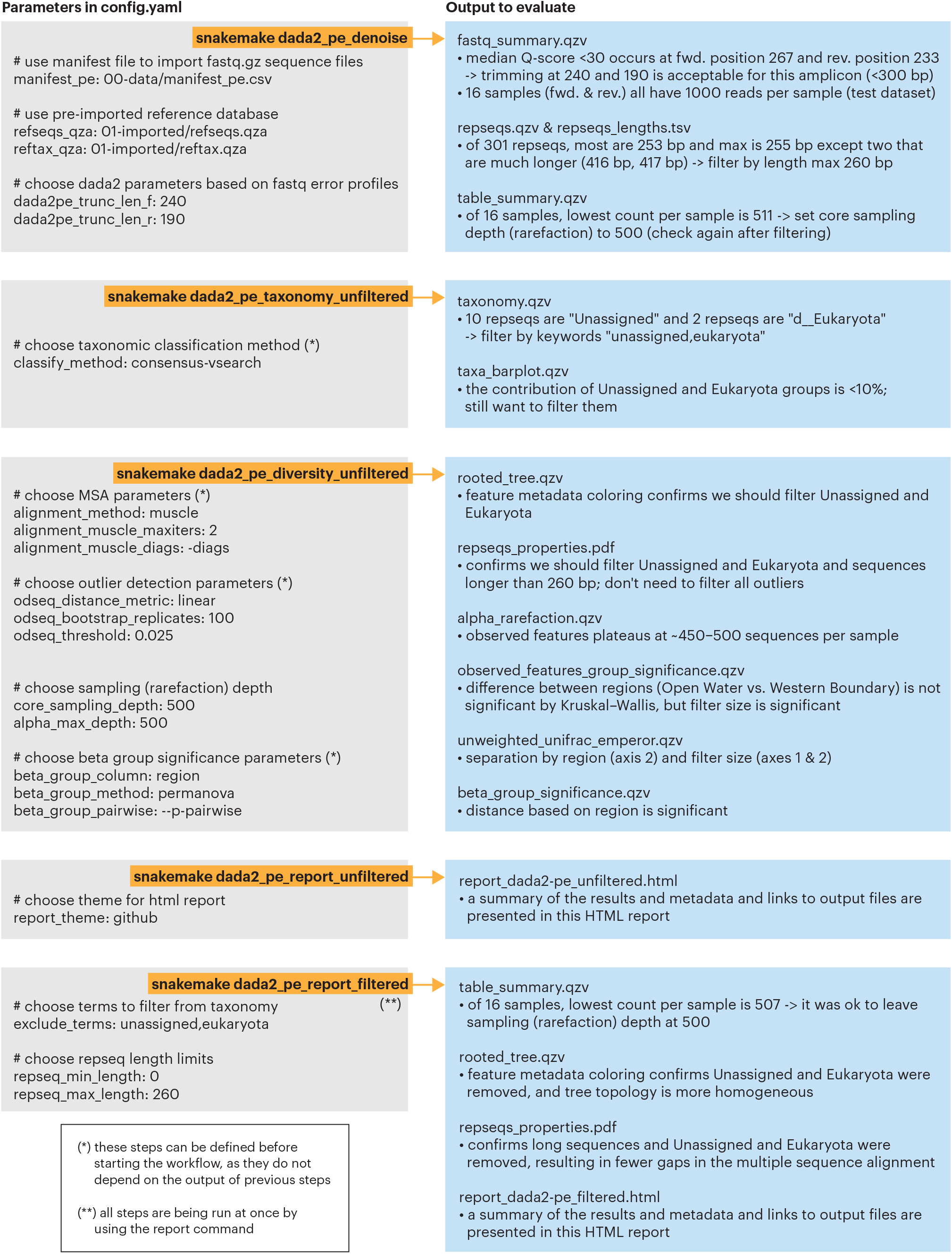
Step-by-step tutorial on Tourmaline using the provided test data, which is subsampled from the 16S rRNA amplicon data of a 2018 survey of Western Lake Erie. Key parameters in *config.yaml* and primary output for each command (pseudo-rule) are listed. Indicated output should be evaluated to determine the appropriate parameters for the next command. Evaluation of the primary outputs and rationale for parameter choice is shown for the test Lake Erie 16S rRNA data that comes with the Tourmaline repository. See Fig. S3 for screenshots of the primary output files.

**Figure 3.**
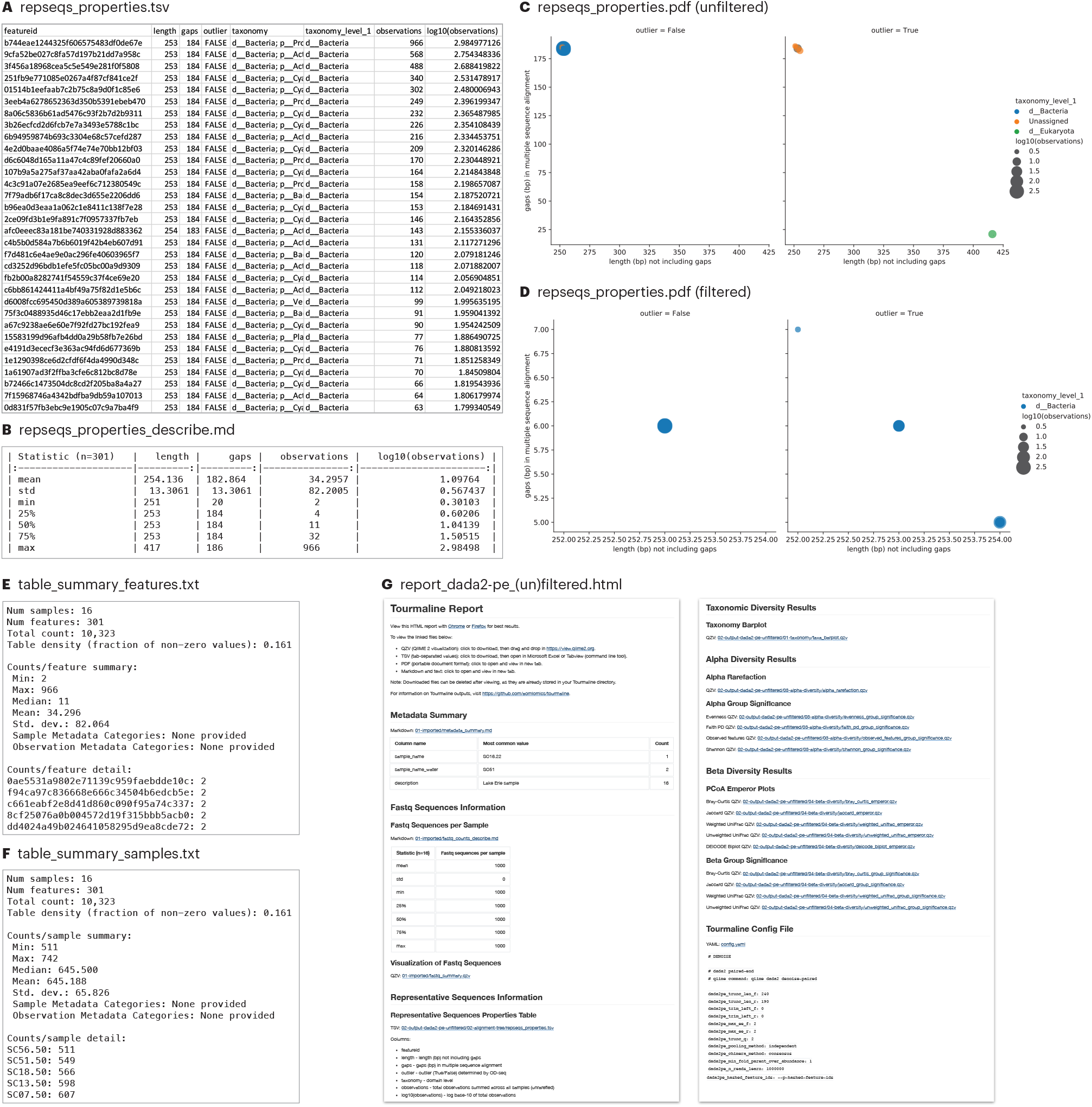
Example of the main outputs of the Tourmaline workflow beyond the QIIME 2 outputs. Contents in panels **A, E, F,** and **G** are truncated. Screenshots of additional output files are provided in Fig. S3. See Fig. 2 for commands(parameters’and guidance.

#### Denoise

The first command is *snakemakedada2_pe_denoise* (Fig. 2), which imports the FASTQ files and reference database (if not already present in directory *01-imported*), summarizes the FASTQ data, runs denoising using DADA2, and summarizes the output. In addition to QIIME 2 visualizations of the feature table, representative sequences, and phylogenetic tree, Tourmaline generates a table and scatter plot (*repseqs_properties.tsv*, *repseqs_properties_describe.md*, and *repseqs_properties.pdf;* Fig. 3A–D) of representative sequence properties, including sequence length, number of gaps in the multiple sequence alignment, outlier status, taxonomy, and total number of observations in the observation table. QC can be performed using *fastq_summary.qzv* (Fig. S3A) for quality scores and *reqseqs.qzv* (Fig. S3C) or *repseqs_lengths.tsv* for representative sequence lengths. The helper script *fastqc_per_base_sequence_quality_dropoff.py* can be run on the output of FastQC and MultiQC to estimate and set DADA2 or Deblur truncation lengths (see below) and then rerun the *denoise* step. Based on the representative sequence lengths, filtering by sequence length can also be set, to be used later in the *filtered* commands. Choice of appropriate sampling (rarefaction) depths for the parameters ‘alpha_max_depth’ and ‘core_sampling_depth,’ to be used in the *diversity* step, can be done by examining *table_summary_features.txt* (Fig. 3E), *table_summary_samples.txt* (Fig. 3F) and *table_summary.qzv* (Fig. S3B).

#### Taxonomy

The second command is *snakemake dada2_pe_taxonomy_unfiltered* (Fig. 2), which assigns taxonomy to the representative sequences using a naive Bayes classifier or consensus BLAST or VSEARCH method and generates an interactive taxonomy table and an interactive barplot of sample taxonomic composition. Choice of taxonomic groups to be filtered by keyword, to be used later with *filtered* commands, can be done by examining *taxonomy.qzv* (Fig. S3D) and *taxa_barplot.qzv* (Fig. S3E).

#### Diversity

The third command is *snakemake dada2_pe_diversity_unfiltered* (Fig. 2), which aligns representative sequences using one of three methods, computes outliers using odseq (Jehl, Sievers and Higgins 2015), and builds a phylogenetic tree. This step generates lists of representative sequences that have unassigned taxonomy and were computed to be outliers, summarizes and plots the representative sequence properties, performs alpha rarefaction, and runs alpha diversity and beta diversity analyses and group significance tests using a suite of metrics. Filtering parameters can be checked by examining *rooted_tree.qzv* (Fig. S3F) and *repseqs_properties.pdf* (Fig. S3G), if desired. Whether sampling depth was sufficient can be evaluated with *alpha_rarefaction.qzv* (Fig. S3I). Alpha and beta diversity patterns and statistically significant differences between groups can be evaluated with *observed_features_group_significance.qzv* (Fig. S3J; other alpha diversity metrics are also provided), *unweighted_unifrac_emperor.qzv* (Fig. S3H; other beta diversity metrics are also provided), and *beta_group_significance.qzv* (Fig. S3K).

#### Report

The fourth and final command is *snakemake dada2_pe_report_unfiltered* (Fig. 2), which creates a comprehensive HTML report of parameters, metadata, inputs, outputs, and visualizations in a single file. The file *report_dada2-pe_unfiltered.html* (Fig. 3G) can be viewed in a web browser, and the linked output files can be viewed in a browser or downloaded and opened with view.qiime2.org (.qzv files) or Microsoft Excel (.tsv files).

#### Filtering

After reviewing the *unfiltered* results—the taxonomy summary and taxa barplot, the representative sequence summary plot and table, and the list of unassigned and potential outlier representative sequences—the user may wish to filter (remove) certain representative sequences by taxonomic group or other properties. This is done by setting the filtering parameters in *config.yaml* and providing a list of any individual representative sequences to filter, then running the *filtered* commands of the workflow: *snakemake dada2_pe_taxonomy_filtered*, *snakemake dada2_pe_diversity_filtered*, and *snakemake dada2_pe_report_filtered* (Fig. 2). Among the *filtered* output, the user can check *table_summary.qzv* (Fig. S3L) to ensure that the sampling depth after filtering did not exclude samples, and examine *rooted_tree.qzv* (Fig. S3N) and *repseqs_properties.pdf* (Fig. S3O) to check that the desired representative sequences were filtered. All of the outputs can be viewed by opening *report_dada2-pe_filtered.html* (Fig. S3M) in a web browser.

### Downstream analysis & meta-analysis

For users who wish to analyze their output further using Jupyter notebooks, we provide Python and R notebooks (github.com/aomlomics/tourmaline/tree/master/notebooks) pre-loaded with popular data analysis and visualization tools for those platforms. These notebooks come ready to run with Tourmaline output, using relative paths to take advantage of Tourmaline’s defined output file structure. The notebooks are shown with the tutorial dataset that comes with Tourmaline. We also provide a Python notebook for meta-analysis, containing commands to merge outputs from multiple Tourmaline runs and then perform diversity analyses on the merged files.

#### Python Jupyter notebook

The Python Jupyter notebook (Fig. S2A) uses the QIIME 2 Visualization and Artifact object classes, loading Visualization and Artifact objects from the .qzv and .qza Tourmaline output files. Before running the notebook, the denoising method, filtering mode, and alpha and beta diversity metrics to be used can be specified by changing variable assignments at the beginning of the notebook. The notebook renders Visualization objects for the feature table summary, representative sequences summary, phylogenetic tree, taxonomy, taxa bar plot, alpha diversity group significance, and beta diversity principal coordinates analysis (PCoA) Emperor plot. Artifact objects can be viewed as a Pandas (McKinney 2010) DataFrame or Series. The notebook generates Pandas DataFrames for the feature table, taxonomy, reference sequence properties, and metadata, and a Pandas Series for alpha diversity. Static plots are generated from some of these tables using Seaborn (Qalieh *et al*. 2017).

#### R Jupyter notebook

The R Jupyter notebook (Fig. S2B) imports Tourmaline artifact (.qza) files using qiime2R (Bisanz 2018) and uses common R packages for analyzing and visualizing amplicon sequencing data, including phyloseq (Halfvarson *et al*. 2017b), tidyverse (Wickham *et al*. 2019), and vegan (Oksanen *et al*. 2020). The notebook covers how to import QIIME 2 count and taxonomy artifact files from Tourmaline into an R environment, merge and manipulate the resulting data frames into a single phyloseq object, and estimate and plot diversity metrics and taxonomy bar plots of the 16S community using phyloseq and other packages. As with the Python notebook, a set of variables can be specified at the beginning of the R notebook to define specific denoising, filtering, and diversity metrics. After reading in the metadata file and merging to a phyloseq object, we define plotting parameters that can be easily modified by the user to customize the R visualizations.

#### Meta-analysis notebook

The meta-analysis notebook (Fig. S2C) guides the user through running Tourmaline on two separate datasets, merging the outputs (feature tables, representative sequences, and taxonomies) and metadata, and performing some basic diversity analyses on the merged output. For simplicity, the two datasets are derived from the test data that comes with Tourmaline. The commands provided could be applied to any set of Tourmaline outputs that the user wishes to combine in a meta-analysis. The only requirement is that the sequenced region must be the same across the datasets for the results to make sense. This notebook is a simple example that demonstrates Tourmaline’s capacity to facilitate merging of outputs and meta-analysis. Many additional analyses are possible on the merged output, such as demonstrated in published microbiome meta-analyses (Thompson *et al*. 2017; Delgado-Baquerizo *et al*. 2018).

### Helper scripts & parameter optimization

Tourmaline comes with several helper scripts that are run automatically with the workflow or run directly by the user. See the Wiki section *Setup* for more information.

#### Initialize a new Tourmaline directory

From the main directory of a newly cloned Tourmaline directory, the script *initialize_dir_from_existing_tourmaline_dir.sh* will copy *config.yaml* and *Snakefile* from an existing tourmaline directory, remove the test files, then copy the data files and symlinks from the existing Tourmaline directory. This is useful when performing a new analysis on the same dataset. The user can clone a new copy of Tourmaline, run this script to copy everything from the old copy to the new one, then make desired changes to the parameters.

#### Create a FASTQ manifest file

Two scripts help create the manifest file that points Tourmaline to the FASTQ sequence files. (1) *create_manifest_from_fastq_directory.py* creates a FASTQ manifest file from a directory of FASTQ files. (2) *match_manifest_to_metadata.py* takes an existing FASTQ manifest file and generates two new manifest files (paired-end and single-end) corresponding to the samples in the provided metadata file.

#### Determine optimal truncation length

If FastQC and MultiQC have been run for Read 1 and Read 2, *fastqc_per_base_sequence_quality_dropoff.py* will determine the position where median per-base sequence quality drops below some fraction (default: 0.90) of its maximum value. This is useful for defining 3’ truncation positions in DADA2 and Deblur (‘dada2pe_trunc_len_f,’ ‘dada2se_trunc_len,’ and ‘deblur_trim_length’).

#### Parameter optimization

The helper scripts and Tourmaline’s standard directory structure enable testing and comparison of different parameter sets to optimize a workflow. By making multiple copies of the directory and populating settings with *initialize_dir_from_existing_tourmaline_dir.sh* script, varying one or a small number of parameters, and running the workflow multiple times in parallel, outputs can be compared visually or programmatically to see the effects of parameter choices and choose a final set. To illustrate this, we analyzed the full dataset of the 2018 Lake Erie 16S rRNA study (BioProject PRJNA679730). Running *fastqc_per_base_sequence_quality_dropoff.py* had suggested that a forward truncation length of 240 bp and reverse truncation length of 190 bp would strike a balance between sequence length and quality, but we wanted to test a full range of truncation lengths. We tested the effects of varying the forward and reverse truncation lengths from 100 bp to 250 bp in 50-bp increments on the distribution of representative sequence length (Fig. S4A) and the number of reads assigned to Eukaryota (Fig. S4B), a group potentially amplified by these primers but with longer representative sequences. This analysis helped choose a set of truncation lengths that would capture a large diversity of target organisms.

### Parallelization & benchmarks

A typical amplicon sequencing dataset is much larger than the test dataset that comes with the Tourmaline repository and will take considerably longer to process. To evaluate runtimes with a real-world dataset, we ran Tourmaline on the full dataset of the 2018 Lake Erie 16S rRNA study (BioProject PRJNA679730), which is the dataset from which the test dataset was subsampled. This dataset was sequenced with 2×300-bp Illumina MiSeq sequencing and consists of 96 samples having an average of 120,338 paired reads per sample, for a total of 11,552,448 paired reads. Processing was performed using the Tourmaline Docker container running on a 2017 iMac Pro with an 18-core 2.3-GHz Intel Xeon W processor and 64 GB RAM (32 GB RAM allocated for the Docker container). Speed improvements with parallelization were tested by running Snakemake with either 1 or 8 cores (parameter: *--cores*). Each main step in the workflow (*denoise, taxonomy, diversity*, and *report*; *unfiltered* commands) was run and timed separately. Times would be expected to be similar for *filtered* commands except that the *denoise* rule does not need to be rerun. The results (Table 1) show that a relatively large dataset of ~100 samples with ~100,000 sequences per sample can be processed with a single core in ~5 hours. Dramatic speed improvements are possible with multiple cores, with this same dataset being processed in ~2 hours when 8 cores were used.

**Table 1.**
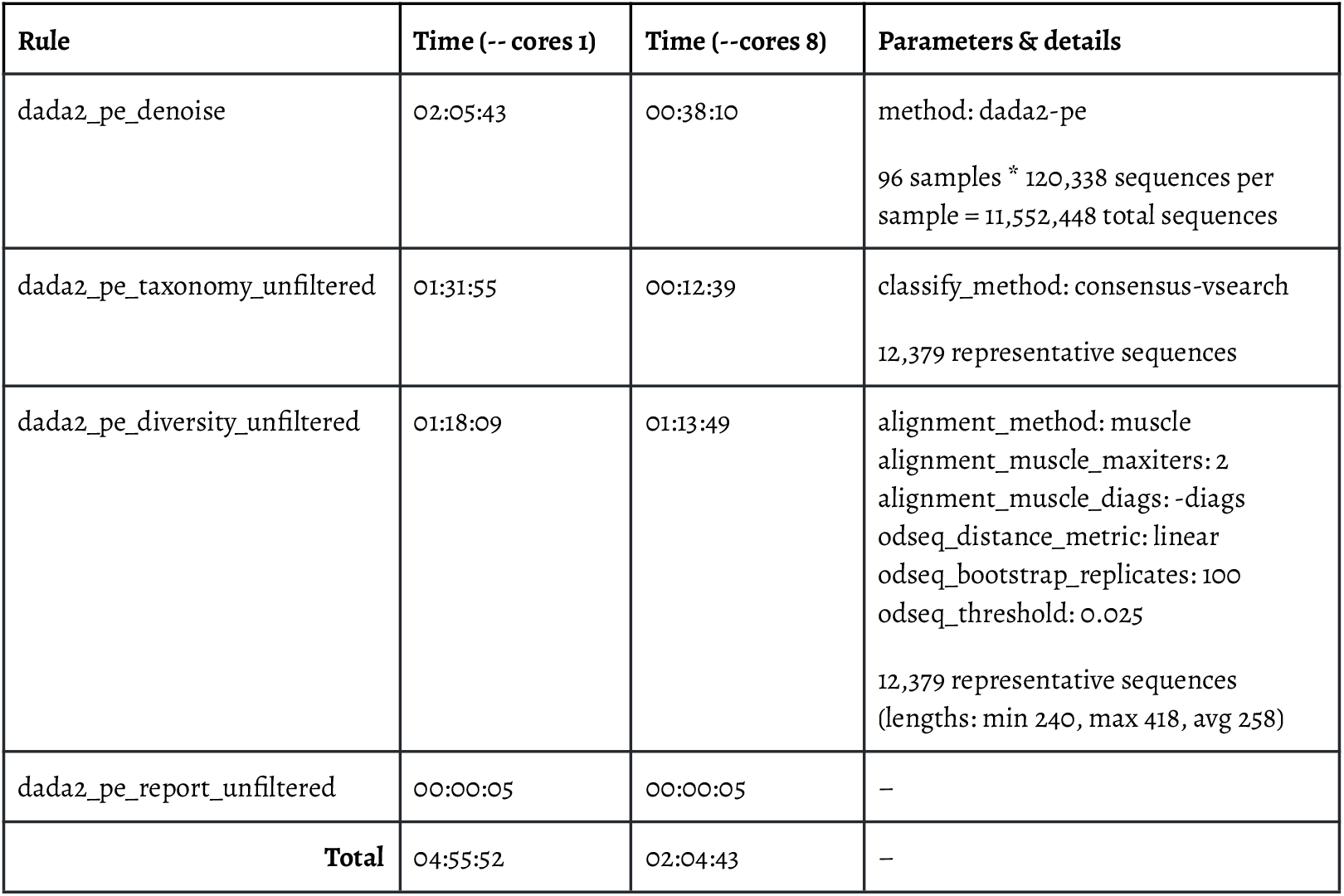
Benchmarking and parallel processing results from running the full 2018 Lake Erie 16S rRNA dataset through Tourmaline with either 1 or 8 cores using a Tourmaline Docker container allocated with 32 GB RAM running on an 18-core iMac Pro (2017). The Snakemake command used the parameter *--cores 1* or *--cores 8*, and parameters in *config.yaml* specifying the number of threads for individual rules were set to 1 or 8, respectively. Times reported are the elapsed real time between invocation and termination and are reported as HH:MM:SS. Times do not include the initial step of importing FASTQ files into a QIIME 2 archive (*fastq_pe.qza*), which took ~2 minutes. Parameters shown in the l**a** st column are those most relevant to the runtimes. Unless otherwise noted, the parameters used were the defaults in *config.yaml*.

### Biological insights

The purpose of performing amplicon sequencing or metabarcoding is to reveal patterns of diversity, community structure, and biological (or environmental) drivers within diverse ecosystems. Whether the system of study is microbial communities in an environmental or biomedical setting or trace environmental DNA in an aquatic or terrestrial system, the kinds of biological questions being asked are similar. Tourmaline supports biological insight in two important ways: (1) by supporting the most popular analysis tools and packages in use today, with capacity to expand as new tools are developed; (2) by providing multiple ways to view the output, giving everyone from experts to novices a platform to visualize and query the output.

Through its core QIIME 2 functionality and downstream support for R and Python data science packages, Tourmaline enables analysis of the core metrics of microbial and eDNA diversity: taxonomic composition, within-sample diversity (alpha diversity), and between-sample diversity (beta diversity). Examining our analysis of the tutorial dataset (Fig. S3), we can see how Tourmaline facilitates insight into Western Lake Erie microbial communities. The interactive barplot (Fig. S3E) provides rapid insights: the most abundant bacterial families in the 5.0-μm fraction are Sporichthyaceae and SAR11 Clade III; the most abundant bacterial family in the 0.22-μm fraction is Cyanobiaceae (the toxic cyanobacterial family Microcystaceae is less abundant), with the largest component assigned as chloroplasts, which can be filtered in a subsequent run; at the domain level, a small fraction of unassigned and Eukaryota-assigned sequences are observed, which can also be filtered. The alpha diversity results show that the 5.0-μm fraction has greater within-sample diversity (number of observed features) than the 0.22-μm fraction (Fig. S3J) and that this diversity appears to be saturated, with a relatively small sampling depth of ~350 sequences per sample sufficient to observe these values (Fig. S3I). However, because a large fraction of the 0.22-μm sequences were identified as chloroplast, filtering out those sequences in a future run would be warranted and provide more accurate diversity results. The beta diversity results show that 16S communities are distinguished both by location (Open Water vs. Western Boundary) and size fraction (0.22-μm vs. 5.0-μm) (Fig. S3H). From this simple tutorial dataset, we demonstrate the use of Tourmaline to analyze environmental amplicon data, in this case revealing the importance of pore size when filtering water samples for microbial sequencing and the presence of spatial variability (regardless of pore size) among microbial communities in Lake Erie.

The ability to view Tourmaline output files with multiple interfaces provides access to researchers with different backgrounds. For users experienced with the Unix command line, the diverse output file types, organized in a defined directory structure, can be queried and analyzed using a wide array of data science tools; anything that can be done with QIIME 2 output and other common sequence diversity output files types can be done with Tourmaline output. For data scientists most comfortable with Jupyter notebooks, the prebuilt Python and R notebooks come ready to work with Tourmaline output and rapidly enable biological discovery from amplicon data. For casual users, the web-based report and QIIME 2 visualizations provide a user-friendly onramp to view and interact with the data. This last mode of interacting with the output opens up amplicon analysis to a wider range of users than is typically possible, from collaborators to students to anyone with limited data science expertise. This increased accessibility can accelerate the pace of discovery by increasing the diversity of researchers able to work with the data.

### Conclusions

Tourmaline provides a comprehensive platform for amplicon sequence analysis that enables rapid and iterable processing and inference of microbiome and eDNA metabarcoding data. It has multiple features that enhance usability and interoperability:

- **Portability.** Native support for Linux and macOS in addition to Docker containers, enabling it to run on desktop, cluster, and cloud computing platforms.
- **QIIME 2.** The core commands of Tourmaline, including the DADA2 and Deblur packages, are all commands of QIIME 2, one of the most popular amplicon sequence analysis software tools available. Users can print all of the QIIME 2 and other shell commands of a workflow before or while running the workflow.
- **Snakemake.** Managing the workflow with Snakemake provides several benefits:
  **– Configuration file** contains all parameters in one file, so the user can see what the workflow is doing and make changes for a subsequent run.
  **– Directory structure** is the same for every Tourmaline run, so the user always knows where outputs are.
  **– On-demand commands** mean that only the commands required for output files not yet generated are run, saving time and computation when re-running part of a workflow.
- **Parameter optimization.** The configuration file and standard directory structure make it simple to test and compare different parameter sets to optimize a workflow.
- **Visualizations and reports.** Every Tourmaline run produces an HTML report containing a summary of metadata and outputs, with links to web-viewable QIIME 2 visualization files.
- **Downstream analysis.** Analyze the output of single or multiple Tourmaline runs programmatically, with qiime2R in R or the QIIME 2 Artifact API in Python, using the provided R and Python notebooks or other code.

Through its streamlined workflow and broad functionality, Tourmaline enables rapid response and biological discovery in any system where amplicon sequencing is applied, from biomedical and environmental microbiology to eDNA for fisheries and protected or invasive species. The QIIME 2-based interactive visualizations it generates allow users to quickly compare differences between samples and groups of samples in their taxonomic composition, within-sample diversity (alpha diversity), and between-sample diversity (beta diversity), which are core metrics of microbial and eDNA diversity. Tourmaline’s unique HTML report and pre-loaded Jupyter notebooks provide ready access to the output, supporting less-experienced researchers and data scientists alike, and the output files are ready to be loaded into a variety of downstream tools in the QIIME 2 and phyloseq ecosystems. Future improvements to the workflow will include support for new QIIME 2 releases and plugins, better integration with Snakemake, possibly including Conda integration and connecting Snakemake’s reporting ability with QIIME 2’s provenance tracking, and enhanced support for cloud computing environments. With its existing features that balance usability, functionality, iterability, and scalability, and with continued development with support from the research community, Tourmaline will be a valuable and longstanding tool for amplicon sequence analysis.

## Methods

### Sample collection and DNA extraction

Water samples were collected using a long-range autonomous underwater vehicle (LRAUV, Monterey Bay Aquarium Research Institute) equipped with a third-generation environmental sample processor (3G-ESP, Monterey Bay Aquarium Research Institute) (Pargett *et al*. 2015). For each sample, water was filtered through stacked 5.0-μm (top) and 0.22-μM (bottom) Durapore filters (EMD Millipore) held in custom 3G-ESP ‘archive’ cartridges and preserved in-cartridge with RNAlater (Thermo Fisher). DNA extraction was performed using the Qiagen DNeasy Blood and Tissue kit.

### Amplicon sequencing

Extracted DNA was amplified using a BiooScientific NEXTFlex 16S V4 Amplicon-Seq Kit 2.0 (NOVA-520999/Custom NOVA-4203-04] (BiooScientific, Austin, TX, USA). Target-specific regions of the forward and reverse primers in the 16S V4 Amplicon-Seq kit were custom ordered to follow the Earth Microbiome Project 16S Illumina Amplicon Protocol: forward primer 515F 5’-GTGYCAGCMGCCGCGGTAA-3’ (Parada, Needham and Fuhrman 2016) and reverse primer 806R 5’-GGACTACNVGGGTWTCTAAT-3’ (Apprill *et al*. 2015). 16S rRNA amplicons were pooled and sequenced on an Illumina MiSeq with 2 x 300-bp chemistry at the University of Michigan Advanced Genomics Core (brcf.medicine.umich.edu/cores/dna-sequencing). Demultiplexed sequences were deposited in NCBI under BioProject PRJNA679730.

## Supporting tables, figures, and videos

Supporting tables and figures are attached to the end of this manuscript.

**Video 1.** Tutorial covering installing and running the Tourmaline workflow, reviewing the report, and viewing QIIME 2 visualization files. https://youtu.be/xKfOxrXBXYQ

## Availability of supporting source code

- Tourmaline is available by cloning the GitHub repository at https://github.com/aomlomics/tourmaline. Tourmaline is released under a 3-clause BSD license.
- Full installation and usage instructions are available from the Tourmaline Wiki at https://github.com/aomlomics/tourmaline/wiki.
- A Docker image is available from DockerHub at https://hub.docker.com/r/aomlomics/tourmaline. This docker image is based on a QIIME 2 Docker image hosted at https://quay.io/repository/qiime2/core?tag=2021.2, and the QIIME 2 license can be found at https://github.com/qiime2/qiime2/blob/master/LICENSE.
- Additional code and data from the full 2018 Lake Erie dataset is available at https://github.com/aomlomics/erie.

## Availability of supporting data

- The test 16S dataset (1000 sequences per sample) is available directly from the GitHub repository at https://github.com/aomlomics/tourmaline.
- Reference databases are available for 16S rRNA at https://docs.qiime2.org/2021.2/data-resources/#silva-16s-18s-rrna and for 18S–ITS rRNA at https://unite.ut.ee/repository.php.
- Output for the tutorial using the included test data are available from Zenodo at https://doi.org/10.5281/zenodo.5044532.
- A snapshot of the GitHub repository will be available from Zenodo upon publication.

## Abbreviations

ASV: amplicon sequence variant
COI: cytochrome oxidase I
DAG: directed acyclic graph
eDNA: environmental DNA
ITS: internal transcribed spacer
NMDS: non-metric dimensional scaling
PCoA: principal coordinates analysis
PCR: polymerase chain reaction
QIIME: Quantitative Insights Into Microbial Ecology

## Author contributions

The Tourmaline workflow was designed and developed by L.R. Thompson. Code was tested by L.R. Thompson, N.V. Patin, S.R. Anderson, and S.J. Lim. The Docker image was built by N.V. Patin and L.R. Thompson. Data analysis and visualization of the case study were done by S.R. Anderson. Analysis notebooks were developed by L.R. Thompson, S.R. Anderson, and G. Sanderson. Samples were collected by P.A. Den Uyl, and K.D. Goodwin. DNA was extracted and prepared for sequencing by P.A. Den Uyl. The manuscript was written by L.R. Thompson, S.R. Anderson, P.A. Den Uyl S.J. Lim, and K.D. Goodwin.

## Acknowledgements

We thank Mehrbod Estaki and Jean Lim for testing of the Tourmaline workflow and feedback on the Tourmaline GitHub repository. We also thank Reagan Errera, Subba Rao Chaganti, Jim Birch, Greg Doucette, for project planning and sample and data collection for the Lake Erie 3G-ESP project. We thank Gregory Dick, Colleen Yancey, and McKenzie Powers for discussions on *Microcystis* diversity and genomics.

---

**Table S1.**
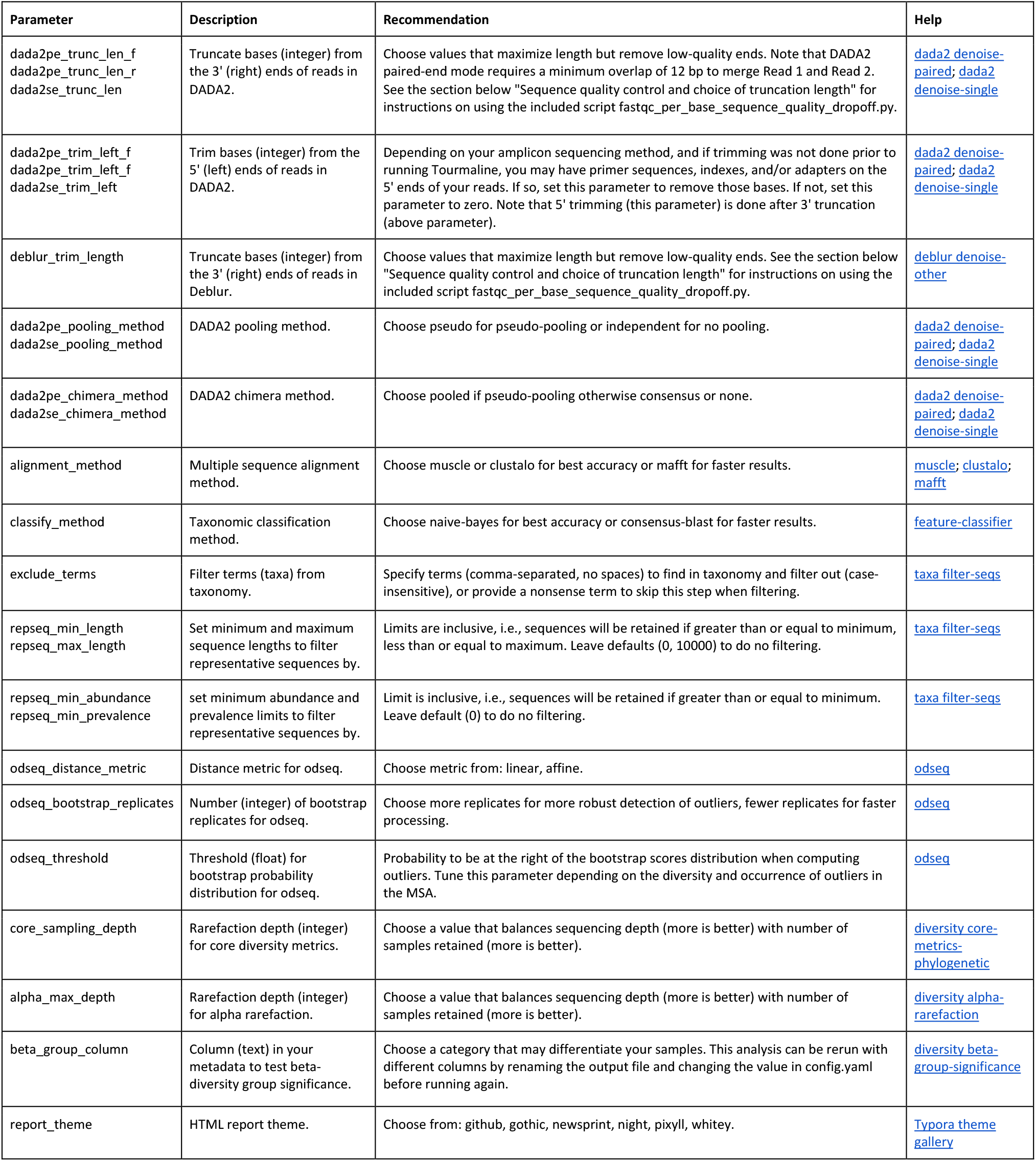
Parameters in the configuration file, *config.yaml*, that the user may edit as necessary. Additional parameters not shown may also be edited. The default configuration file is provided in the top level of the GitHub repository. The file format of *config.yaml*, YAML (yet another markup language), is a simple markup language that is used by Snakemake to specify parameters for a workflow.

**Figure S1.**
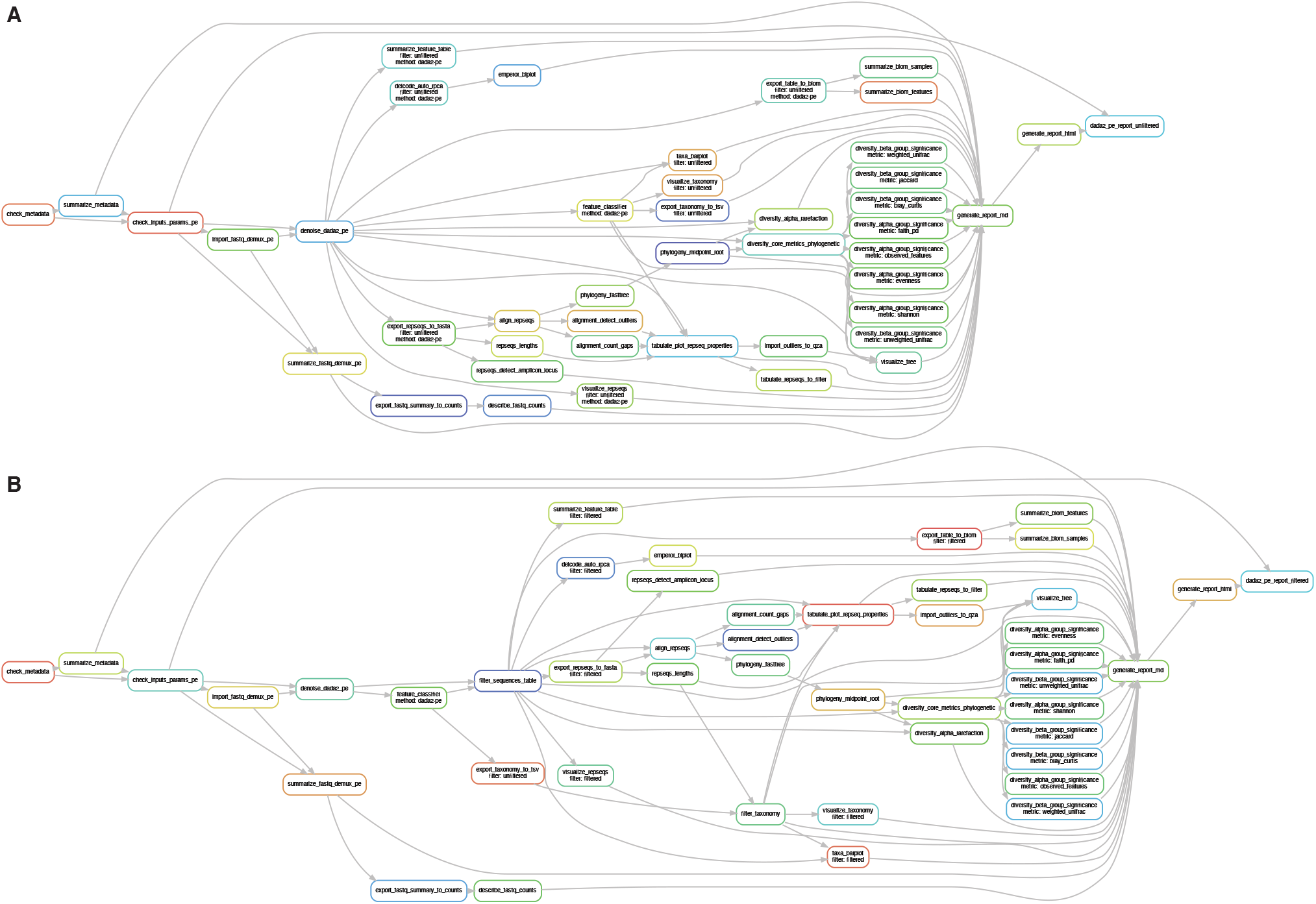
Directed acyclic graphs (DAGs) of the Tourmaline workflow for the DADA2 paired-end method from start to report with (a) *unfiltered* commands and (b) *filtered* commands. This figure was generated from the test data that comes with the repository by running the commands (a) *snakemakedada2_pe_report_unfiltered --dag | dot -Tpdf-Grankdir=LR -Gnodesep=0.1 -Granksep=0.1 >dag_pe_report_unfiltered.pdf* and (b) *snakemakedada2_pe_report_filtered --dag | dot -Tpdf-Grankdir=LR - Gnodesep=0.1 -Granksep=0.1 > dag_pe_report_filtered.pdf*. For a simpler graph, substitute *--rulegraph* for *--dag* in the above commands.

**Figure S2.**
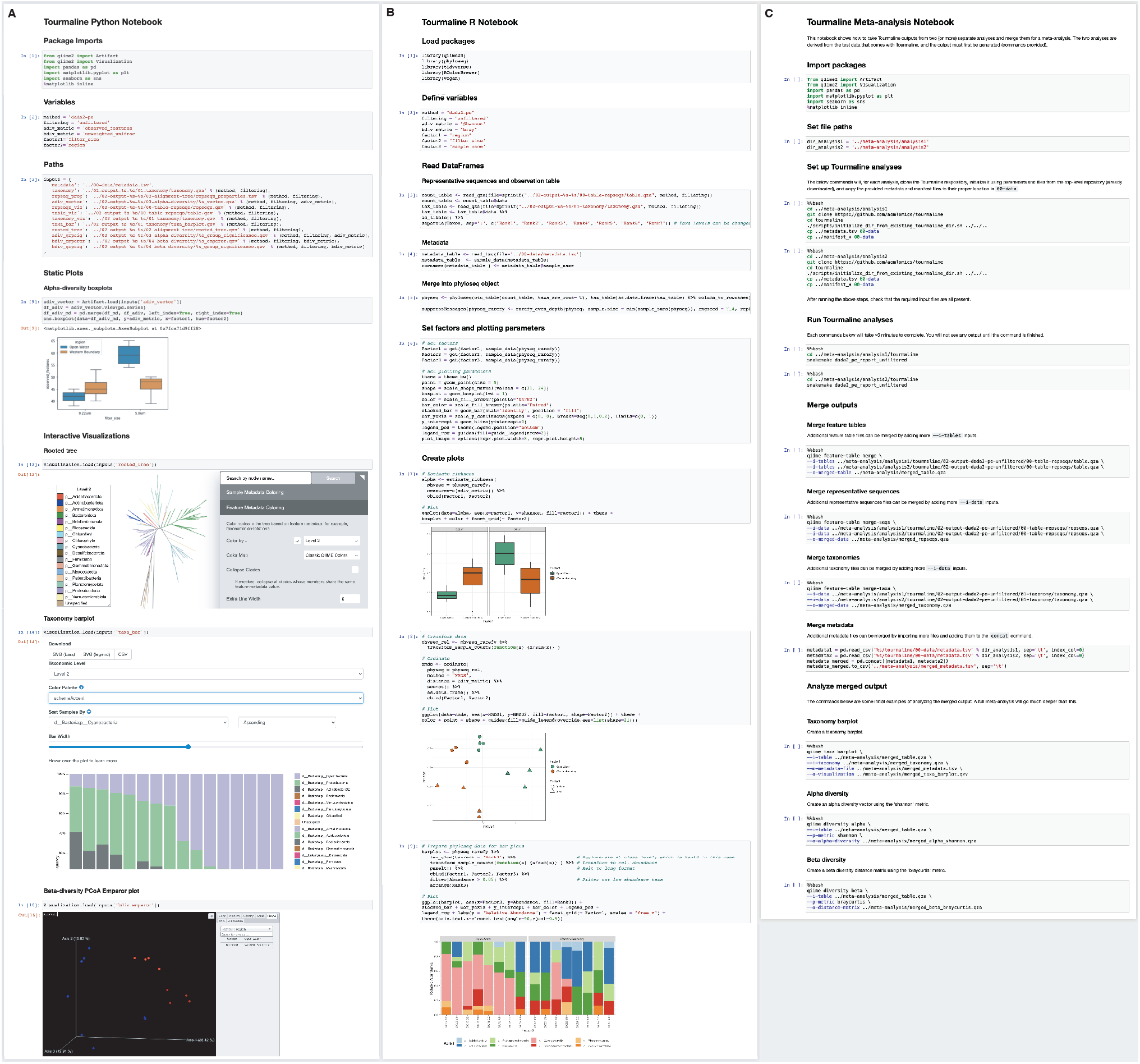
Screenshots of Tourmaline’s included Python and R Jupyter notebooks running the provided test data. Both notebooks are designed to run out-of-the-box with the Tourmaline output from any dataset. (**A**) The Tourmaline Python notebook loads and displays sample metadata, feature metadata (representative sequences properties and taxonomy), static plots generated by Seaborn, and interactive QIIME 2 visualizations. (**B**) The Tourmaline R notebook demonstrates how to load .qza files (counts and taxonomy) into R, merge files with metadata into a single phyloseq object, and generate high-quality visualizations of community diversity and taxonomy using phyloseq and suite of tidyverse packages (e.g., ggplot2). (**C**) The Tourmaline meta-analysis notebook walks through the merging of two sets of Tourmaline outputs and performing some basic diversity analyses on the merged files. The number of processed datasets being merged in the meta-analysis can be increased by adding additional inputs to the commands.

**Figure S3.**
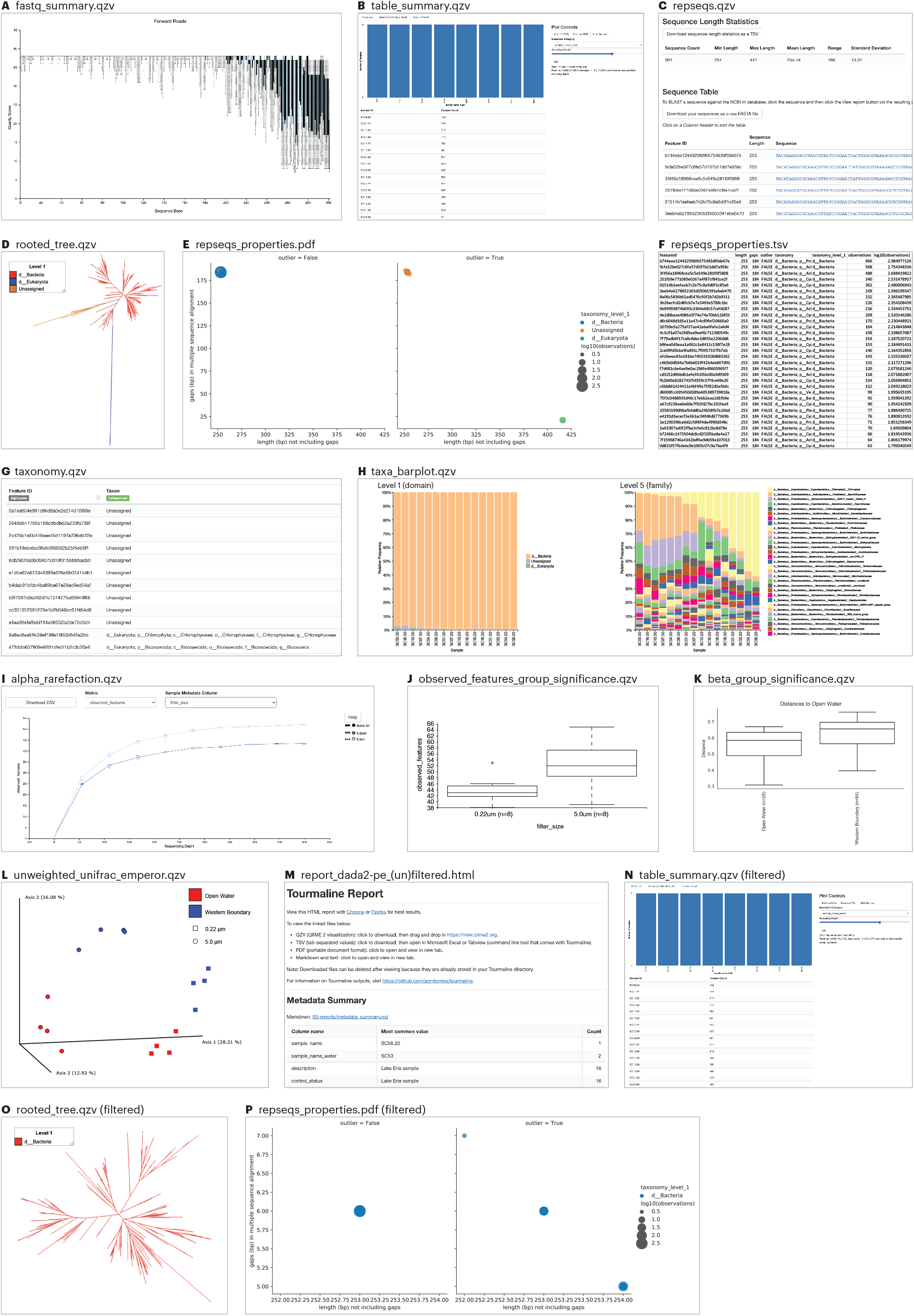
Screenshots of the primary output files after running Tourmaline on the test data (see Fig. 2 for commands, parameters, and guidance). The visualization files (.qzv, .pdf, .html) are useful both for data evaluation and discovery and for biological insight.

**Figure S4.**
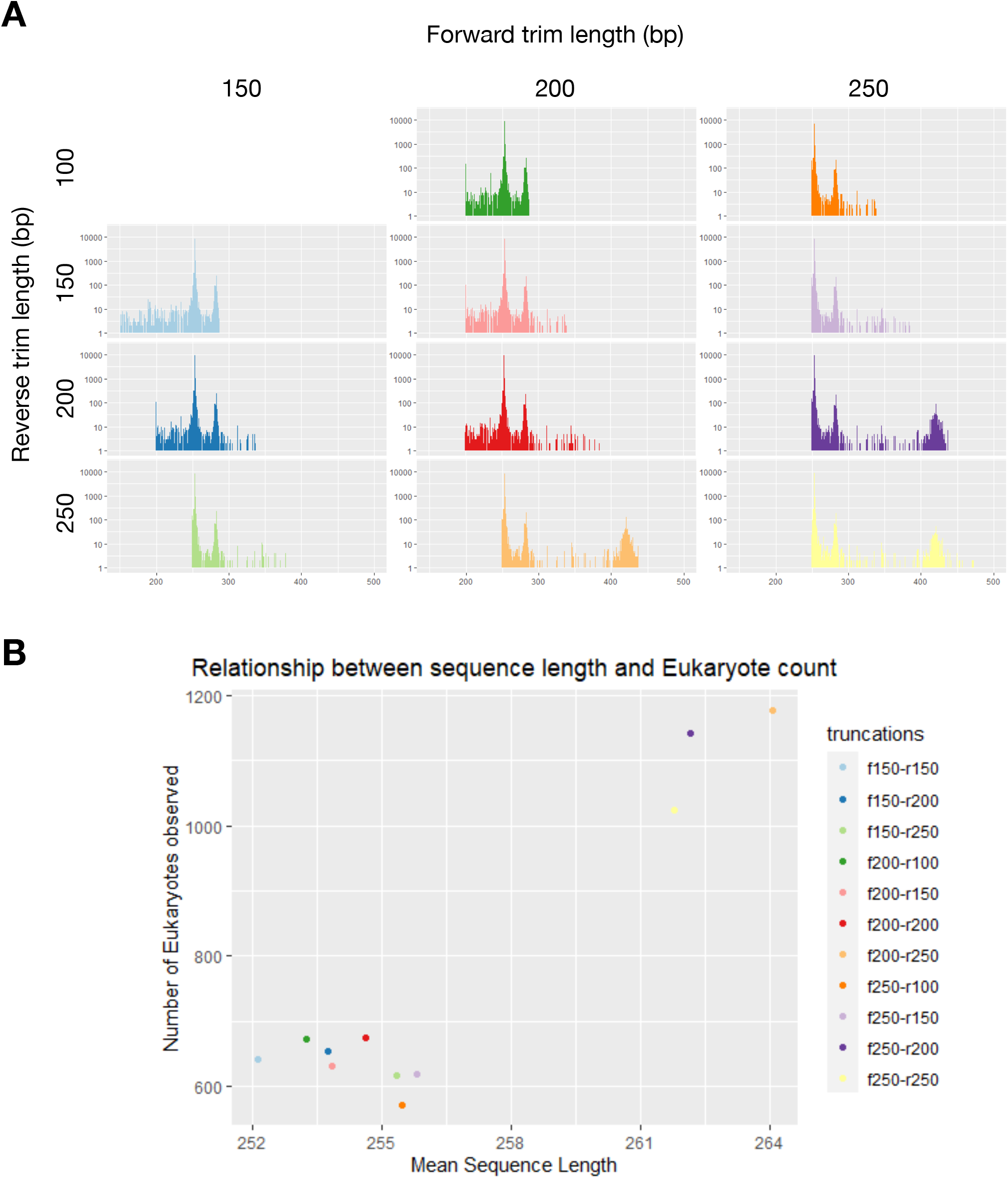
Effect of truncation length parameters on (**A**) the distribution of representative sequence length and (**B**) the number of reads assigned to Eukaryota in the full 2018 Lake Erie 16S rRNA amplicon data.

